# Global prevalence of potentially pathogenic short-tandem repeats in an epilepsy cohort

**DOI:** 10.1101/2020.08.20.259168

**Authors:** Claudia Moreau, Jacques L. Michaud, Fadi F. Hamdan, Joanie Bouchard, Vincent Tremblay, Berge A. Minassian, Patrick Cossette, Simon L. Girard

**Affiliations:** Department of Fundamental Sciences, University of Quebec in Chicoutimi, Chicoutimi, Canada; CHU Sainte-Justine Research Center, Montreal, Quebec, Canada; Department of Neurosciences and Department of Pediatrics, University of Montreal, Montreal, Quebec, Canada; Department of Pediatrics, University of Montreal, Montreal, Quebec, Canada; Department of Pediatrics, Hospital for Sick Children and University of Toronto, Toronto, Canada; Department of Pediatrics, University of Texas Southwestern, Dallas, TX, USA; CHUM Research Center, Montreal, Canada; Department of Neurosciences, University of Montreal, Montreal, Canada

**Keywords:** Short tandem repeats, epilepsy, neurodevelopmental diseases

## Abstract

This study aims to decipher the role of short tandem repeats (STRs) in epilepsy patients. Whole genome short-read sequencing data of 734 epileptic patients was used to look for known STR expansions associated with increased risk of neurodevelopmental diseases or epilepsy using three different software. Results show one hit of particular interest on *ARX* gene associated with Early Infantile Encephalopathic Epilepsy that could be causal for one patient with developmental and epileptic encephalopathy. However, we show that the different software do not agree on most of the calls above the threshold and that experimental validation is still needed for diagnostic, although these algorithms could prove useful for pre-selection of samples to be validated.

## Introduction

Epilepsy affects ~3% of individuals; half of these cases start during childhood. Although monogenic forms of the disease have been reported(1–4), they represent less than 2% of epilepsy cases. In the last decade, many studies have been conducted on different genetic and bio-statistic methods to better understand the complex genetic mechanisms underlying epilepsy(5). Recently, a large GWAS on epilepsy identified 16 loci associated with the disease, many of which were already known or suspected(6). Moreover, copy number variants were studied in the context of epilepsy(7–9). Polygenic risk scores have also been used to try to better understand the complexity of the disease(10,11). Despite these efforts, there is still a substantial missing heritability component in epilepsy genetics(12).

There is growing evidence that short tandem repeats (STRs) affect gene expression(13) and play a role in genetic disorders(14,15) notably in epilepsy(16–20). However, until recently, most STR expansions were not detectable in next-generation short-read sequencing datasets because they exceeded the reads’ length and therefore, were not included in genetic studies(21). Fortunately, this has recently changed and such methods with accompanying software are now available(22–26).

In this study we used TREDPARSE(25), Expansion Hunter(22) and GangSTR(26) on whole genome short-read sequencing data of epileptic patients to look for known STR expansions associated with increased risk of neurodevelopmental diseases or epilepsy.

## Data and methods

This study was approved by the CHUM Research Center (CRCHUM) ethics committee and written informed consent was obtained for all patients.

### Phenotyping of patients

The epilepsy cohort was composed of families with at least three affected individuals with Idiopathic Generalized Epilepsy (IGE), Non-Acquired Focal Epilepsy (NAFE) or Developmental Epileptic Encephalopathy (DEE) previously collected in CHUM Research Center in Montreal and in the Hospital for Sick Children in Toronto as part of the Canadian Epilepsy Network (CENet) and diagnosed by neurologists. The clinical epilepsy phenotype is defined based on the Classification of the Epilepsy Syndromes established by the International League against Epilepsy (ILAE) (27).

More specifically for NAFE, patients were at least five years of age and have experienced at least two unprovoked seizures in the six months prior to starting treatment, an MRI scan of the brain that did not demonstrate any potentially epileptogenic lesions, other than mesial temporal sclerosis.

Patients with clinical and EEG characteristics meeting the 1989 ILAE syndrome definitions for IGE were included. An MRI of the brain was not required for participation. All patients were at least four years of age at the time of diagnosis. In IGE, we also included patients with Jeavons syndrome, which is an idiopathic generalized form of reflex epilepsy characterized by childhood onset, unique seizure manifestations, striking light sensitivity and possible occurrence of generalized tonic-clonic seizures.

The DEE patients were also recruited by the CENet group at three centers: the Sainte-Justine University Hospital Center in Montreal, the Toronto Western Hospital, and the Hospital for Sick Children in Toronto. They included subjects with diverse DEE phenotypes. The criteria used for the selection of these individuals were as follows: (1) intractable epilepsy defined as an absence of response to two appropriate and well-tolerated anti-epileptic therapies (AEDs) over a 6-month period and an average of at least one focal, generalized tonic-clonic, myoclonic, tonic, atonic, or absence seizure or epileptic spasm per month during the period of poor control; (2) ID or global developmental delay; (3) absence of malformations or focal and multifocal structural abnormalities on brain MRI; and (4) absence of parental consanguinity and family history of epilepsy, ID, or autism in first-degree relatives.

In this study we used WGS of 218 IGE from 186 families (between one and three individuals per extended family), 218 NAFE from 198 families (ranging from one to four members) and 298 DEE from 297 families.

### Libraries preparation and whole-genome sequencing

Samples were sequenced at 30X coverage at Genome Quebec Innovation Center in Montreal. gDNA was cleaned using ZR-96 DNA Clean & ConcentratorTM-5 Kit (Zymo) prior to being quantified using the Quant-iTTM PicoGreen dsDNA Assay Kit (Life Technologies) and its integrity assessed on agarose gels. Libraries were generated using the TruSeq DNA PCR-Free Library Preparation Kit (Illumina) according to the manufacturer’s recommendations. Libraries were quantified using the Quant-iTTM PicoGreen dsDNA Assay Kit (Life Technologies) and the Kapa Illumina GA with Revised Primers-SYBR Fast Universal kit (Kapa Biosystems). The average size fragment was determined using a LabChip GX (PerkinElmer) instrument. The libraries were denatured in 0.05N NaOH and diluted to 8pM using HT1 buffer. The clustering was done on an Illumina cBot and the flowcell was run on a HiSeq 2 500 for 2×125 cycles (paired-end mode) using v4 chemistry and following the manufacturer’s instructions. A phiX library was used as a control and mixed with libraries at 0.01 level. The Illumina control software used was HCS 2.2.58 and the real-time analysis program used was RTA v. 1.18.64. bcl2fastq v1.8.4 was used to demultiplex samples and generate fastq reads. The filtered reads were aligned to reference Homo_sapiens assembly b37. Each readset was aligned using BWA-MEM version 0.7.10 to create a Binary Alignment Map file (.bam).

### Analyses

We looked at the number of repeats in patients at 29 trinucleotide-repeat diseases (summary in (21,25)) and six STR expansions previously shown to be associated with an increased risk of epilepsy (table S1) using TREDPARSE(25), Expansion Hunter(22) and GangSTR(26) with default parameters.

We considered hits above the threshold if two out three software called an expansion(28). When this was possible, we genotyped the parents of the patient to validate the call. We additionally simulated reads of known repeats number around the expansion threshold, aligned them using BWA-MEM as described in(26) and applied the algorithms on the resulting bams.

## Results

The distribution of the number of repeats in patients for 29 trinucleotide-repeat diseases associated with neurological conditions is presented in figure 1 and for six STR expansions associated with an increased risk of developing epilepsy is shown in figure 2 for the three software. The filled dots denote alleles above the reported risk threshold. Table 1 shows the number of patients above the risk threshold for the three software. We can clearly see a difference in the number of expansions called by the three algorithms, and that Expansion Hunter called much more expansions than the other two software.

**Figure 1:**
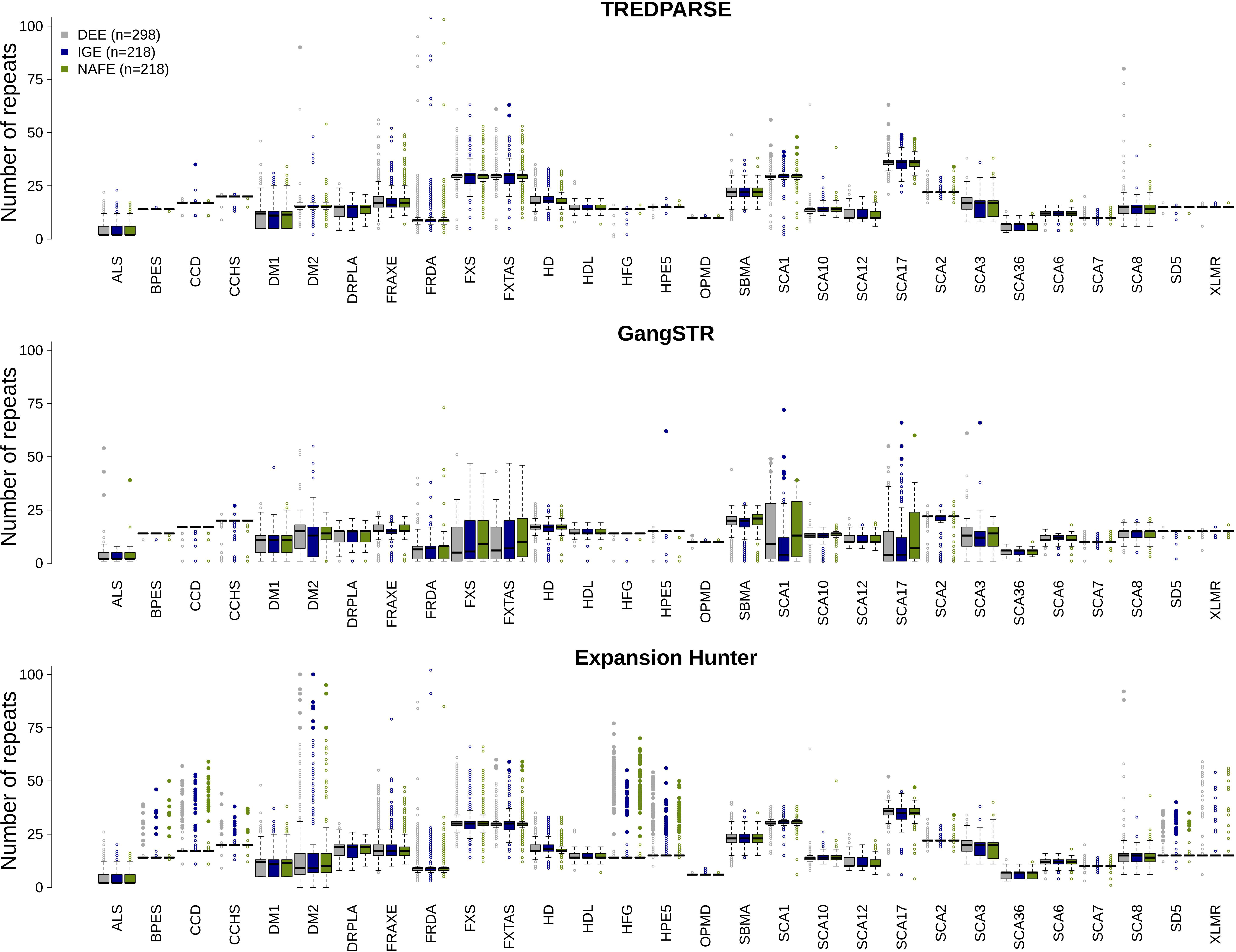
Number of repeats for 29 trinucleotide-repeat diseases for three software. Boxplots of the number of repeats for 29 trinucleotide-repeat diseases for DEE (grey), IGE (blue) and NAFE (green). Filled dots are above the risk threshold. Graphs were cut at 100 repeats for better visibility, note that very few individuals have expansions higher than 100 repeats.

**Figure 2:**
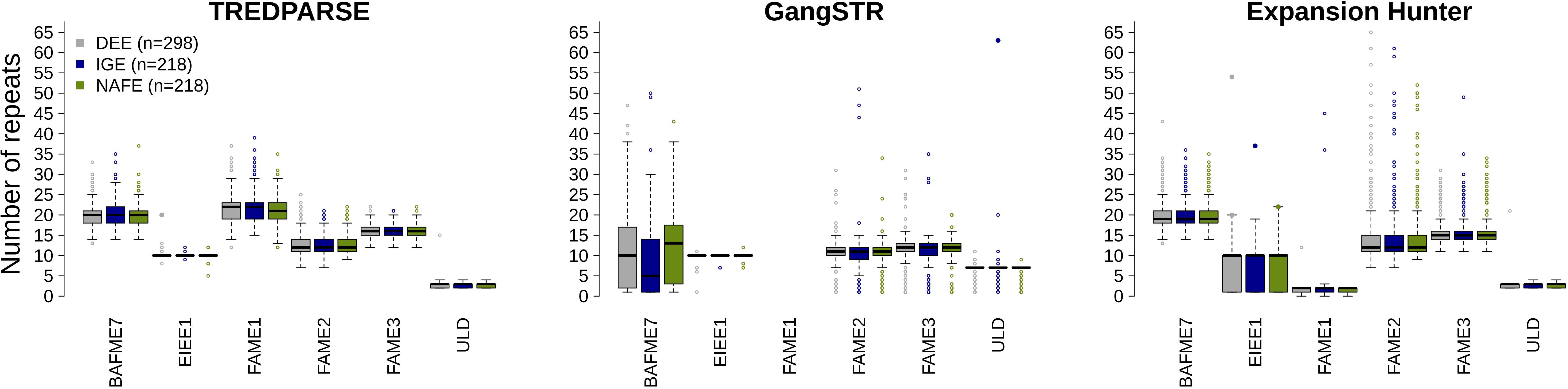
Number of repeats for six epilepsy related STRs for three software. Boxplots of the number of repeats for six epilepsy related STRs for DEE (grey), IGE (blue) and NAFE (green). Filled dots are above the risk threshold. Graphs were cut at 65 repeats for better visibility, note that very few individuals have expansions higher than 65 repeats.

**Table 1 :** Number of individuals above the threshold for 15 repeat expansion diseases and 2 STRs associated with epilepsy.

| **STR** | **Expansion Hunter** | **TREDPARSE** | **GangSTR** |
| --- | --- | --- | --- |
| **ALS** | 0 | 0 | 4 |
| **BPES** | 24 | 0 | 0 |
| **CCD** | 80 | 1 | 0 |
| **CCHS** | 24 | 0 | 1 |
| **DM2** | 32 | 1 | 0 |
| **FXTAS** | 10 | 3 | 0 |
| **HFG** | 181 | 0 | 0 |
| **HPE5** | 95 | 0 | 1 |
| **OPMD** | 0 | 0 | 1 |
| **SCA1** | 0 | 10 | 9 |
| **SCA17** | 2 | 10 | 5 |
| **SCA2** | 1 | 1 | 0 |
| **SCA3** | 0 | 0 | 2 |
| **SCA8** | 3 | 1 | 0 |
| **SD5** | 50 | 0 | 0 |
| **EIEE1** | 4 | 1 | 0 |
| **ULD** | 0 | 0 | 1 |

Table 2 presents the individuals for whom there are at least two software calls above the threshold for one STR. There are only four such STRs, three associated with neurodevelopmental diseases (FXTAS, SCA2, SCA8) and one associated with Early Infantile Encephalopathic Epilepsy (EIEE1, OMIM #308350). Table 2 also compares the carrier frequency of our patients with the 12,632 individuals in the original TREDPARSE study(25) and the reported prevalence when this information was available. The carrier frequency for the four expansions in our cohort is higher than the carrier frequency in the 12,632 individuals previously reported except for FXTAS where our patients’ carrier frequency is lower than the reported carrier frequency, but still higher than TREDPARSE controls. Of note, we report one hemizygote (using TREDPARSE) male DEE being above the risk threshold for the *ARX* gene known for its association with EIEE1 compared to 0 carrier in the TREDPARSE study. We could validate that this STR expansion was transmitted by his (heterozygote) mother using TREDPARSE although the mother call was not validated using Expansion Hunter and the patient was called heterozygote (both alleles above threshold).

**Table 2 :** STRs above threshold for at least two software.

| STR | Phenotype | Sex | inheritance | Risk Threshold | Hg19 repeats | Patient TRED | Patient Exp Hunter | Father TRED | Mother TRED | Father Exp Hunter | Mother Exp Hunter | Present study count (n=734) | TRED controls count<sup>a</sup> (n=12,632) | Reported prevalence<sup>b</sup> (/100,000) |
| --- | --- | --- | --- | --- | --- | --- | --- | --- | --- | --- | --- | --- | --- | --- |
| FXTAS | IGE | M | XLD | 55 | 20 | 44-58 | 55-55 |  |  |  |  | 1 | 2 | *Carrier freq : 300-500* |
| SCA2 | NAFE | F | AD | 33 | 23 | 22-34 | 22-34 |  |  |  |  | 1 | 4 | 1,5 |
| SCA8 | DEE | M | AD | 80 | 15 | 12-80 | 12-92 |  |  |  |  | 1 | 3 | 0,5 |
| EIEE1 | DEE | M | XLR | 20 | 10 | 20-20 | 20-54 | 10-10 | 20-10 | 10-1 | 10-1 | 1 | 0 | NA |

<sup>a</sup> The TREDPARSE numbers of at risk individuals were taken from table 1 (25).
<sup>b</sup> Prevalence (or carrier frequency for FXTAS) was reported along with references in supplementary table S1 (25).

To further look at the accuracy of TREDPARSE and Expansion Hunter calls for four STRs above the threshold, we simulated reads of known repeat numbers (figure 3). The threshold is represented by the blue dashed lines. We can see that there are no false positives for SCA2 and EIEE1 even very close to the threshold.

**Figure 3:**
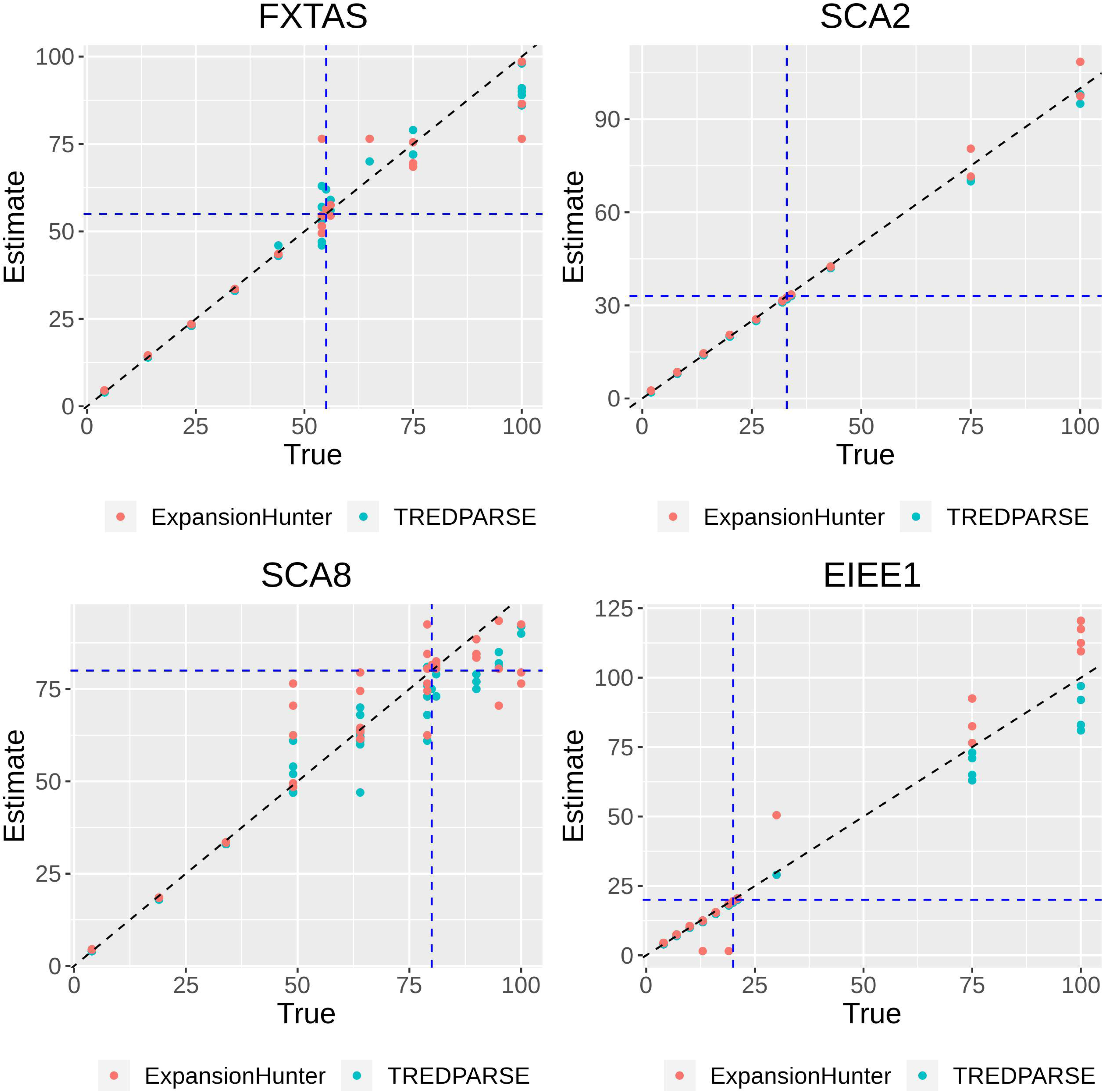
Simulations vs real data for four STR expansions for two software. Real (x axis) vs simulated (y axis) number of repeats for TREDPARSE (cyan) and Expansion Hunter (pink). The diagonal dashed line represents the truth and the blue dash lines the thresholds.

## Discussion

We tested known STR expansion diseases associated with an increased risk of developing neurodevelopmental conditions and epilepsy in 734 epilepsy patients. We found patients above the risk threshold for four out of the 29 STR expansion diseases and the six epilepsy related STR expansions tested. For these, the carrier frequency in our patients is higher than what was reported for 12,632 samples in the study(25) and also higher than the known prevalence or carrier frequency except for FXTAS (table 2). This is interesting since comorbidities and shared genetic mechanisms are frequent between neurological disorders and epilepsies(29,30). Of course, this might also be due to the small sample size in the present study.

Moreover, our patients are from the Quebec population which is a well-documented founder population (31). In fact, some of these expansion diseases have already been documented <u>as</u> being more frequent in particular Quebec regions as a result of the founder effect(s). Of note, OPMD(32), DM1(33) and FRDA(34). For these three, we do not report any patient above the risk threshold with at least two software. However, Eastern Quebec regions are known for successive regional founder events that could explain frequency changes, but our patients are mainly from the Western Quebec area which is not one of those regions with a marked founder effect. Consequently, we do not think that the higher carrier frequency observed in the present study for the STR expansion diseases can be explained by the founder effect.

For the recessive *ARX* gene STR expansion causing EIEE1, we see a hemizygote DEE boy (with his heterozygote mother) who is above the risk threshold confirmed by Expansion Hunter although less convincing. The boy has been diagnosed with DEE in the neonatal period and was successfully treated with clonazepam. His DEE was accompanied by microcephaly and the degenerative process of white matter. The patient was negatively tested for the *STXBP1* gene in single gene testing and we did not find any de novo variant for this patient in a study including 200 DEE trios sequenced for the whole genome(9). Consequently, we believe that the STR expansion in the *ARX* gene is causal.

Nevertheless, the distribution of the number of repeats for the three algorithms does not show a high correlation. There were many calls above the threshold that were not validated by another software as shown in tables 1 and 2. Validations with two out of three software, were found only between Expansion Hunter and TREDPARSE. None of the GangSTR calls was validated by another program. Moreover, the distribution of repeats using Expansion Hunter show a lot of hits above the threshold compared to both other software (figure 1). To further measure the accuracy of TREDPARSE and Expansion Hunter at the four STR sites showing an expansion, we simulated reads of known repeats’ number (figure 3). Expansion Hunter and GangSTR software have previously been reported as very accurate for huge expansions (over 1000 repeats) (Figure S2 in (26)) while TREDPARSE is less accurate above 100-200 repeats. However, the thresholds for most expansions are below 100 repeats. The goal here was to simulate around the threshold to see how the software performs into that repeats’ range. The dashed blue lines represent the thresholds. For SCA2 and EIEE1, the simulations reflect the truth even at one repeat around the threshold, the software did not call false positives. On the other hand, for FXTAS and SCA8, it seems more likely that the expansion calls would in fact be false positives around the threshold.

## Conclusion

In this study, we performed a survey of STRs in epilepsy patients. We found four expansions on STRs previously associated with neurodevelopmental disease or epilepsy. Interestingly, we found one hit on the *ARX* gene that could be causal for one DEE patient. Nevertheless, we note a lack of consensus among the different software used and an uncertainty around the risk threshold. These software could not be used as diagnostic tools, but for some STRs such as SCA2 and EIEE1, they could prove useful as pre-diagnostic tools to select cases for further experimental validation.

## Data availability statement

Raw whole genome sequences of a subset of the epilepsy patients for which we have appropriate consent have been deposited in the European Genome-phenome Archive, under the accession code EGAS00001002825. The STR data calculated using the three software will be available upon request.

## Acknowledgments

We are thankful to Compute Canada/Calcul Québec for the access to storage and computing resources. We are extremely grateful to all patients and their families for participating in this research.

## Supplemental Material

**Table S1:** Description of STRs previously identified in epilepsy disorders.

