## Supplemental Table 1 for "Global prevalence of potentially pathogenic short-tandem repeats in an epilepsy cohort"

**Table S1 : Description of STRs prviously identified in epilepsy disorders.**

| Chr | Start | Stop | Gene | Reference motif | Disease | Hg19 Ref | Risk threshold |
|---|---|---|---|---|---|---|---|
| chr2 | 96862805 | 96862859 | *STARD7* | AAAAT | FAME2 | 11 | 340 |
| chr4 | 160263679 | 160263768 | *RAPGEF2* | TTTTA | BAFME7 | 18 | 60 |
| chr5 | 10356451 | 10356519 | *MARCH6* | TTTTA | FAME3 | 12 | 791 |
| chr8 | 119379052 | 119379357 | *SAMD12* | TTTCA[a] | FAME1 | 0 | 440 |
| chr21 | 45196326 | 45196360 | *CSTB* | CGCGGG GCGGGG | ULD | 3 | 30 |
| chrX | 25031779 | 25031808 | *ARX* | CGC | EIEE1 | 10 | 20 |

If the number of repeats of the expansion was not available, we inferred it using the minimum length of this expansion reported in the literature.

[a] This motif is not seen in the reference. Reference is TTTTA
